# Haplotype assembly of autotetraploid potato using integer linear programming

**DOI:** 10.1101/346429

**Authors:** Enrico Siragusa, Richard Finkers, Laxmi Parida

## Abstract

Haplotype assembly of polyploids is an open issue in plant genomics. Recent experimental studies on highly heterozygous autotetraploid potato have shown that available methods are not delivering satisfying results in practice. We propose an optimal method to assemble haplotypes of highly heterozygous polyploids from Illumina short sequencing reads. Our method is based on a generalization of the existing minimum fragment removal (MFR) model to the polyploid case and on new integer linear programs (ILPs) to reconstruct optimal haplotypes. We validate our methods experimentally by means of a combined evaluation on simulated and real data based on 83 previously sequenced autotetraploid potato cultivars. Results on simulated data show that our methods produce highly accurate haplotype assemblies, while results on real data confirm a sensible improvement over the state of the art. Binaries for Linux are available at: http://github.com/ComputationalGenomics/HaplotypeAssembler.

## 1 Introduction

Many plants of agronomic importance are polyploids, that is their somatic cells contain more than two copies (*p* > 2) of each haploid set of chromosomes. Seedless varieties of watermelon and banana are human-induced triploids (*p* = 3); cultivars of potato, peanut, cotton, tobacco and coffee are naturally occurring tetraploids (*p* = 4); wheat is hexaploid (*p* = 6). Knowledge of the haplotypes, the distinct sequences of haploid chromosomes, is limited or absent even for common cultivars. This fact limits the effectiveness of plant breeding to selectively develop particular phenotypic traits.

An economical way of determining haplotypes is bulk DNA sequencing followed by haplotype assembly. The haplotypes are sequenced jointly and then demultiplexed *in silico* by assembling sequenced DNA fragments based on a known reference haplotype. A study by Uitdewilligen *et al.* [15] demonstrated the genotyping by targeted resequencing of a large collection of autotetraploid potato cultivars using Illumina HiSeq instruments. In that context, assembling haplotypes from standard Illumina paired-end data should be feasible because potato is highly heterozygous (median density of heterozygous SNVs is around 70 bp, depending on target region). Despite that, a subsequent experimental study by [10] reported that existing computational methods for haplotype assembly are not delivering satisfying results in practice. It is unclear to what extent assemblies are inaccurate because of heuristics, or insufficient or erroneous data.

### Previous work

Haplotype assembly of diploid genomes has been extensively studied over the past seventeen years. Lancia *et al.* [8] first introduced the problem and proposed a combinatorial model called *minimum fragment removal* (MFR) that is solvable in polynomial time for contiguously sequenced fragments (i.e., single-end reads) but is NP-hard for gapped fragments (i.e., reads obtained via paired-end or mate-pairs protocols). Subsequently, Lippert *et al.* [9] refined MFR as *minimum error correction* (MEC), which is NP-hard even for contiguous fragments. MEC became the de-facto model to assemble diploid genomes as several exact and heuristic methods have been proposed for that. We refer the reader to [12] for a comprehensive treatment of haplotype assembly for diploids.

In recent years, the focus shifted towards assembling polyploid genomes. Aguiar and Istrail [1] defined an NP-hard problem named *minimum weighted edge removal* in a compass graph and employed a minimum-cost spanning tree heuristic to solve it. Berger *et al.* [3] defined a probabilistic framework and used heuristic branch-and-bound to find likely haplotypes given the fragments. Das and Vikalo [5] casted the problem as a correlation clustering problem and derived approximate solutions to the associated semi-definite program via lagrangian relaxation followed by randomized projections and greedy refinement. Xie *et al.* [17] defined a NP-hard problem called *polyploid balanced optimal partition* and proposed constrained dynamic programming to find heuristic solutions. The combinatorial problem for polyploids is particularly hard and heuristic methods are delivering suboptimal solutions.

Integer linear programming (ILP) is a powerful mathematical programming method to efficiently solve combinatorial problems to optimality [16]. Schwartz *et al.* [12] remarked that “the ILP strategy has thus so far received comparatively little attention in the haplotype assembly field”. Chen *et al.* [4] proposed an ILP that is however specific to diploids. In a different setting, Szolek *et al.* [14] successfully applied an MFR-based ILP to haplotype human leukocyte antigen (HLA) genes from short-sequencing reads. To the best of our knowledge, haplotype assembly for polyploids has not been attacked yet using ILP.

### Our contribution

We propose an optimal method to assemble haplotypes in highly heterozygous polyploids from Illumina short sequencing reads. Our method is based on a generalization of MFR to the polyploid case; we leverage ILP to reconstruct optimal haplotypes. In addition, we propose haplotyping distance as a general method to perform pairwise comparison of polyploids and we apply that to assess the accuracy of haplotype assemblies. We validate our models experimentally through a combined evaluation on simulated and real autotetraploid potato sequencing data extrapolated from the targeted sequencing of 83 cultivars performed by Uitdewilligen *et al.* [15]. Results on simulated data show that MFR-based ILPs achieve mean 98 % haplotyping recall and precision, that is a 4–11 % improvement over existing tools. Results on real data confirm the superiority of our methods in terms of genotyping and read error correction.

## 2 Methods

### 2.1 Haplotype assembly

#### Problem definition

Let us fix 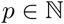 as organism ploidy and 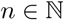 as number of observed loci in a genomic region of interest. Such genomic region consists of *p* latent haplotypes *H* = {*h*_1_*,…,h_p_*} over the *n* genomic loci. At each locus *j* we observe 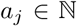 distinct alleles across all haplotypes in *H*. We univocally encode alleles as integers so that each latent haplotype *h_k_* is a sequence of alleles *h_k_*_1_*…h_kn_* with *h_kj_* ∊ [0*, a_j_*). Figure 1a shows an example of latent haplotypes.

**Fig. 1:**
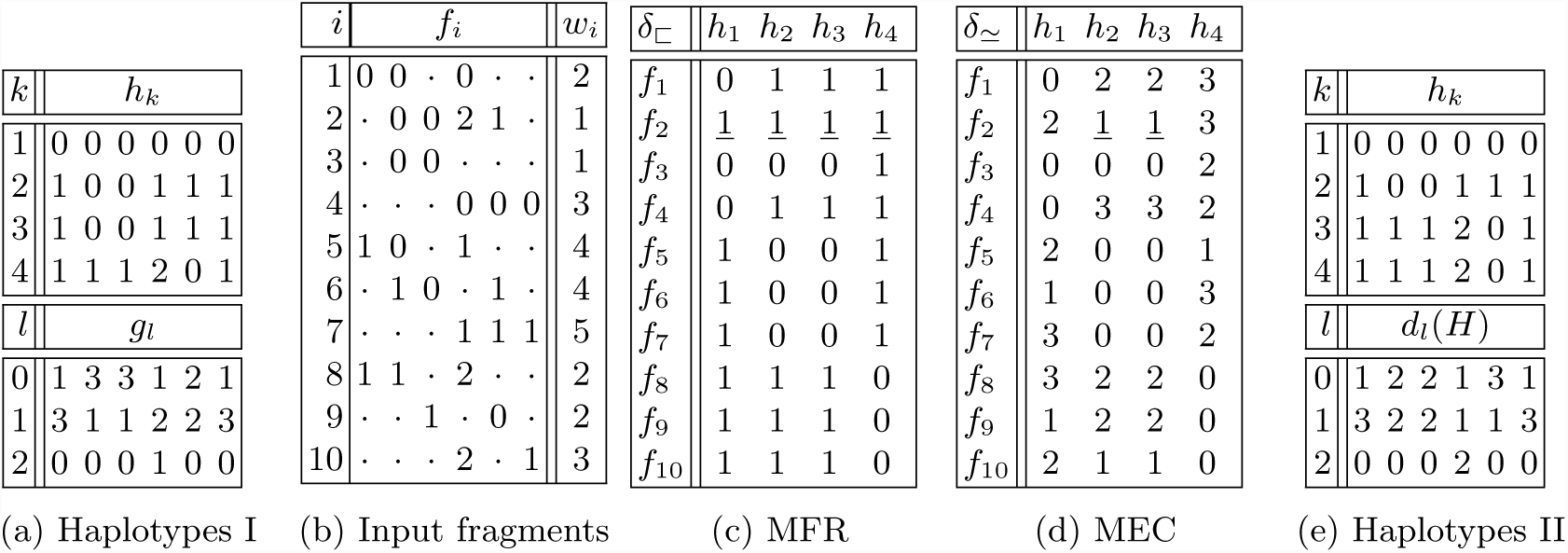
Toy example of haplotype assembly. Table (a) shows a set of latent haplotypes *H* along with their genotypes. Table (b) shows a multiset (*F, W*) of fragments coming from *H*. Table (c) shows values of *δ*_⊏_ between *F* and *H*. Objective function FR (*F, W*; *H*) has value 1 as fragment *f*_2_ with multiplicity *w*_2_ = 1 is not contained in any haplotype. Table (d) shows function *δ*_≃_. Likewise, objective function EC has value 1. Table (e) shows an alternative assembly of the fragments. Both latent and assembled haplotypes yield MEC and MFR 1. Assembled haplotypes in (e) become infeasible when known genotype constraints *G* are incorporated into the MFR or MEC model.

We are given in input a multiset (*F, W*) of *preprocessed* fragments coming from all the latent haplotypes in *H*, where fragments in *F* = {*f*_1_*,…,f_m_*} have multiplicities in *W* = {*w*_1_,…,*w_m_*}. Each fragment *f_i_* is a gapped sequence of alleles *f_i_*_1_*…f_in_* with *f_ij_* ∊ [0*, a_j_*) ∪ {·} and symbol · denoting unknown alleles, i.e., missing observations due to paired sequencing protocols or quality thresholds. Fragments may contain sequencing errors. The haplotype assembly problem for polyploids is to find the *p* latent haplotypes in *H* given all observations from fragments in (*F, W*). Figure 1b illustrates an example of input fragments.

#### Heterozygous loci

In haplotype assembly we ignore *homozygous* loci, i.e., genomic loci with only one observed allele. At homozygous loci either all latent haplotypes have the same observed allele or we have no information whether observations are wrong or missing. For this reason, we perform haplotype assembly only on *heterozygous* loci, i.e., genomic loci for which *a_j_* ≥ 1.

#### Concordance and containment

Before introducing combinatorial models for haplotype assembly, we define basic relations of concordance and containment between sequences of alleles. Let *a* and *b* be two allele observations at the same genomic locus. Given *a* ≠ · and *b* ≠ ·, we say that *a* and *b* are concordant if *a* = *b* and discordant if *a* ≠ *b*. If either *a* = · or *b* = ·, we say that *a* and *b* are non-discordant and write *a* ≃ *b*. In addition, if *b* ≠ · and either *a* = · or *a* = *b*, we say that *a* is contained in *b* and write *a* ⊏ *b*. Consequently, we say that a sequence of alleles *x* is contained in *y* if all alleles of *x* are contained in those of *y* (and the same for concordance). Two non-discordant fragments contribute to explain a common latent haplotype, while two discordant fragments must either originate from distinct haplotypes or imply some sequencing error.

#### MFR and MEC for polyploids

Two combinatorial models called minimum fragment removal (MFR) [8] and minimim error correction (MEC) [9] have been proposed for diploids. These models are motivated by the parsimony principle by which, within a set of possible explanations of observations, the simplest one is most likely to be true. In this instance, the closest possible haplotypes to input fragments, according to some predefined distance function, are most likely to be correct. Here we generalize MFR and MEC to polyploids.

We first define an objective function FR that counts the number of erroneous fragments that are not contained in candidate haplotypes *H* and must be therefore removed from (*F, W*) to explain a correct assembly of *H*:

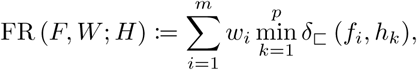

where function *δ*_⊏_ (*f_i_, h_k_*) is defined as 0 if *f_i_* ⊏ *h_k_* and 1 otherwise. Consequently, MFR is computed by finding the haplotypes *H*^∗^ that minimize the number of fragments to be removed:

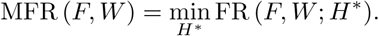

Analogously, we define MEC as the sum of minimum Hamming distances between haplotypes *H* and input fragments (*F, W*). MEC is based on the following objective function:

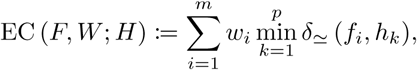

where function *δ*_≃_ (*f_i_, h_k_*) denotes the Hamming distance between *f_i_* and *h_k_* parameterized by ≃. Hence EC counts the minimum number of base calling errors in (*F, W*) to explain a correct assembly of *H*.

Figures 1c–1d illustrate tabulated values for objective functions FR and EC. Note how a fragment *f_i_* can be non-discordant with (or have equal distance to) more than one haplotype *h_k_*. We also remark that our generalization of MFR and MEC do not partition *F* in *k* disjoint subsets as done for instance by Das and Vikalo [5] or Xie *et al.* [17].

Motivated by the low error-rate and high coverage of Illumina high-throughput sequencing data, we develop our haplotype assembly model on MFR rather than MEC. The simplicity of MFR let us formulate concise and efficient ILPs.

#### Genotype-based MFR

MFR alone is not well-defined in the polyploid case (neither MEC is). Figures 1a and 1e illustrate this issue. A notion of coverage is necessary to determine the number of copies of each unique haplotype, while these simple combinatorial models are clearly coverage oblivious. For this reason we supplement MFR with genotyping information, that is essentially a surrogate of coverage information. Genotypes are estimated *a priori* from fragments (*F, W*) assuming uniform sequencing coverage across the haplotypes. MEC-based models can be easily extended in the same way.

Our extended model takes as additional input a matrix of genotypes *G*, where 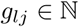 denotes the dosage (i.e., multiplicity) of the *l*-th allele observed at locus *j* and 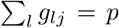. We use this additional information to narrow down the search space of our model. Our genotype-constrained MFR model (*c*MFR) determines the latent haplotypes as:

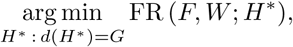

where *d*(*H*) denotes the genotypes matrix induced by candidate haplotypes *H*. Note how this is the natural generalization of the *all-heterozygous assumption* for diploids by which all observed heterozygous alleles are believed to be correct and the two latent haplotypes must be complementary.

Alternatively, we incorporate genotyping information in the objective function as the *L*^1^ norm between induced and input genotypes. Our genotype-augmented MFR model (*a*MFR) determines the latent haplotypes as:

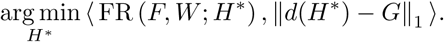

In this alternative model, we prioritize MFR over genotyping information. That is useful whenever sequencing coverage is too low to call genotypes accurately.

#### ILP for genotype-constrained MFR (cMFR)

Our ILPs are based on containment relation ⊏ between fragments and haplotypes. In what follows, unless otherwise stated, subscript *i* ∊ [1*, m*], *j* ∊ [1,*n*], *k* ∊ [1,*p*]. We define three types of binary variables to encode respectively fragments to be removed (Line 2), fragments contained in haplotypes (Line 3) and alleles on each haplotype (Line 4). Variable *x_i_* is 0 if fragment *f_i_* is to be removed and 1 otherwise. Variable *y_ik_* is 1 if *f_i_* ⊏ *h_k_* and 0 otherwise. Variable *z_kjl_* is 1 if haplotype *h_k_* has the *l*-th allele at locus *j* and 0 otherwise. For convenience, our ILP maximizes the inverse of function FR and *y_ik_* tabulates function *δ*_⊏_ in negated form with respect to Figure 1c.

Objective function 1 maximizes the number of unremoved fragments in *R* weighted by multiplicities in *W*. Constraint 5 marks a fragment for removal unless it is contained in some haplotype. Constraints 6–7 allow a fragment to be contained in a haplotype only if all its alleles are contained in the haplotype. Constraint 8 improves convergence by observing that if *f_i_* ⊏ *f_o_* and 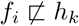 then 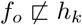. Finally, Constraint 9 imposes input genotype information on the haplotypes.
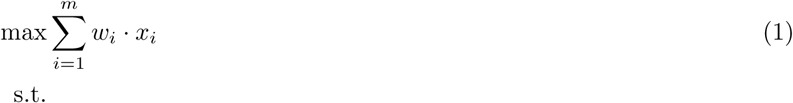

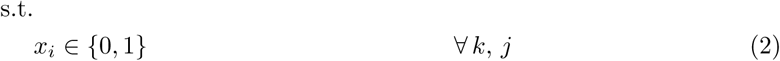

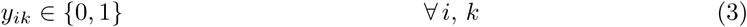

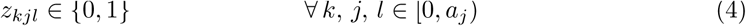

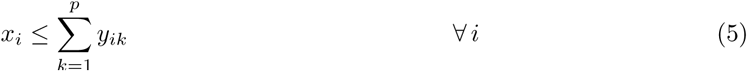

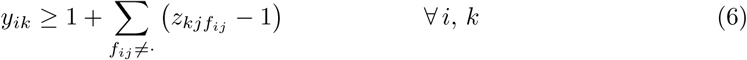

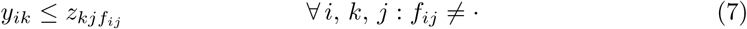

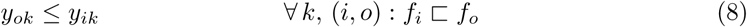

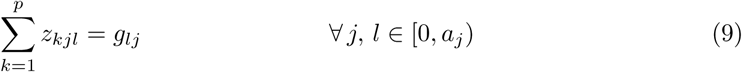

The above ILP may become infeasible when some haplotypes lack sequencing coverage locally. If an haplotype has no coverage at locus *j* with allele *l*, there will be strictly less than *g_lj_* haplotypes with allele *l*. To consider this case, we substitute Constraint 9 with 11–12 (and keep Constraints 2–8). Now we have to minimize also the distance between input and induced genotypes, that is:

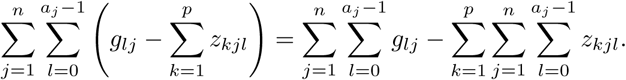

We drop the first term because it is a constant (*np*) and scale the second term by the sum of fragment multiplicities in order to prioritize the genotypes over MFR. This leads us to Objective function 10. 
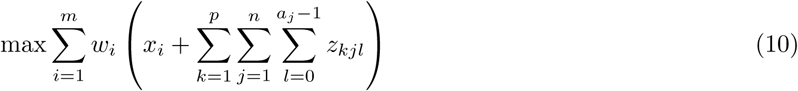

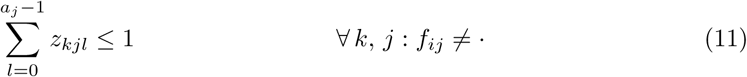

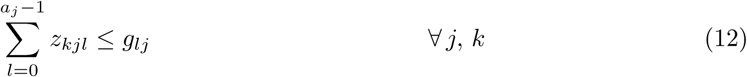

#### ILP for genotype-augmented MFR (aMFR)

Genotyping is subject to errors in regions of low coverage or in presence of allelic bias introduced during fragment selection or amplification prior to sequencing. For this reason, we propose an alternative MFR model that is not hardly constrained by genotypes and gives precedence to fragments. We keep Constraints 2–8 from the previous ILPs and introduce real variables *d_jl_* (Line 14) to model absolute distances between input and induced genotypes. We substitute Constraint 12 with 15–16 and incorporate absolute distances in Objective function 13.
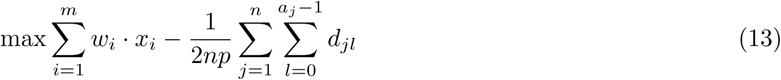

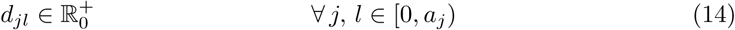

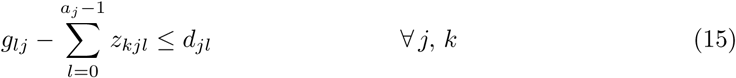

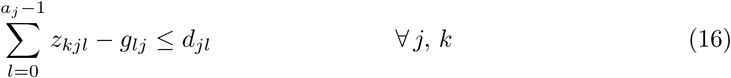

### 2.2 Haplotype comparison

We now describe methods to compare two polyploids *X*, *Y* over a common genomic region. In particular, we propose *haplotyping distance* as a general method for pairwise comparison of polyploids with arbitrary genotypes. For simplicity, in what follows we assume that all alleles in *X* and *Y* are known, i.e., there is no ·.

#### Phasing distance

We call phasing distance the minimal Hamming distance between haplotypes in *X* and any permutation of the haplotypes in *Y*:

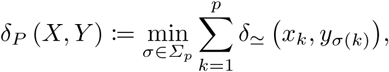

where *Σ_p_* denotes the set of all permutations of sequence {1, 2,…,*p*} and *σ*(*k*) the element at position *k* in permutation *σ*.

We compute phasing distance in 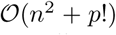 time by proceeding in two easy steps. First, we compute the Hamming distance between all pairs of haplotypes in *X* and *Y*. Second, we permute haplotypes in *Y* to minimize the sum of Hamming distances with respect to the ordered haplotypes in *X*.

#### Haplotyping distance

Phasing distance does not model haplotype *switch* operations. Therefore, in terms of mismatches, a switch between two pairs of corresponding haplotypes can be more or less costly depending on its genomic position. A switch towards the center of the genomic region is accounted as a long sequence of mismatches rather than a single operation with unitary cost. Figures 2a–2c show examples of switches.

**Fig. 2:**
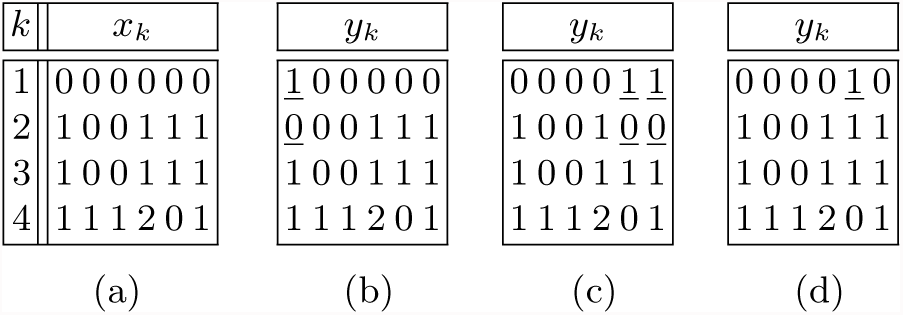
Toy example of haplotype comparison. Phasing distance between (a) and (b) is 2, while it is 4 between (a) and (c). On the contrary, vector and haplotyping distance is 2 in both cases (note how suffixes of haplotypes *y*_1_ and *y*_2_ can be switched in c). Distance between (a) and (d) is undefined in vector error, while it is 1 in phasing and haplotyping distance.

Berger *et al.* [3] introduced *vector error* as a generalization of *switch error* to count the number of switches between two polyploid genomes. Vector error has limited applicability on practical instances because it is defined only for genomic regions with equal genotypes (see Figure 2d). To overcome this limitation, we propose *haplotyping distance* as a generalization of vector error.

Haplotyping distance counts the minimum number of haplotype switches and genotype mismatches between two polyploids. To compute haplotyping distance we extend the dynamic programming algorithm for vector error given by Xie *et al.* [17]. Algorithm 1 operates on *X* and *Y* transposed, i.e., it advances one column at time from left to right. Initialization (Line 3) counts the number of mismatching alleles at column *j* = 1 between *X* and each permutation *σ* ∊ *Σ_p_* of the haplotypes in *Y*. Recurrence (Line 6) minimizes haplotyping distance for each column *j >* 1 building up from distance at column *j* − 1 plus the minimum distance obtained by permuting the haplotypes in *Y* as *τ* ∊ *Σ_p_* and paying for mismatching alleles as well as switches introduced by going from permutation *σ* to *τ*. Algorithm 1 computes haplotyping distance in 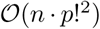 time and 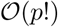 space.

##### Algorithm 1

Dynamic programming calculation of haplotyping distance. Note that the algorithm works columnwise from left to right, i.e., it operates on the transposed haplotypes.

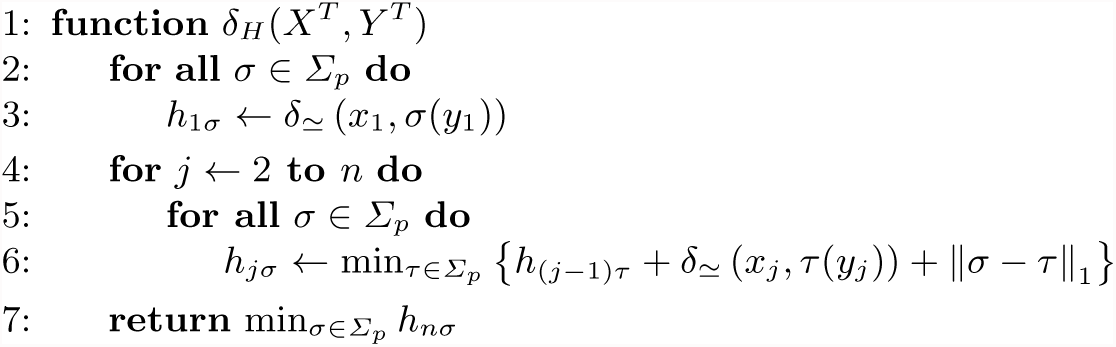

## 3 Results

### Real data

We obtained targeted high-throughput sequencing data for 83 highly heterozygous autotetraploid (*p* = 4) potato cultivars by Uitdewilligen *et al.* [15]. Cultivars have been sequenced on Illumina HiSeq 2000 at 63× median coverage with 2 × 96 *bp* paired-end reads from fragments of 300 *bp* mean length and 60 *bp* standard deviation. We reproduced the analysis pipeline described in [15] for read mapping, deduplication, variant calling and genotyping. We produced a total of 996 test instances by selecting 12 high-quality genomic regions from the sequencing panel. Table S1 shows the genomic coordinates of the selected regions, as well as the median number of heterozygous variants per sample.

### Simulated data

We simulated tetraploid data matching the real genotypes of the 83 potato cultivars by Uitdewilligen *et al.* [15]. First, we used the haplotype simulator SimBA-hap [13] to generate 80 samples from each genomic region with genotypes fitting those of the sequenced cultivars. We remark that simulated variants and genotypes are biallelic and error-free. Subsequently, we used the read simulator Mason [6] to produce Illumina-like sequencing reads from each simulated region. We produced an Illumina-like paired-end dataset reflecting the technical specifications of actual Illumina instruments: 90× coverage with 2 × 150 *bp* paired-end reads from fragments of 500 *bp* mean length and 60 *bp* standard deviation at 0.3 % mean sequencing error. To assess the effect of sequencing errors and fragments length on haplotype assembly, we simulated two supplementary datasets by altering specific simulation parameters. We produced a second paired-end dataset consisting of error-free paired-end reads. Subsequently, we simulated a mate-pair dataset (with sequencing errors) using fragments of 1000 *bp* mean length and 400 *bp* standard deviation. We produced a total of 960 test instances per dataset.

### Infrastructure

We implemented our ILPs for MFR in C++ using IBM CPLEX^®^ 12.7.0 and the software library SeqAn 2.3.2 [11]. In all experiments we configured CPLEX timeout at 600 seconds. We ran and evaluated our MFR-based models against HapCompass [1], SDhaP [5], H-PoP [17] on all real and simulated instances. While HapCompass accepted standard BAM and VCF files, for SDhaP and H-PoP we had to produce intermediate fragment files using scripts provided by Motazedi *et al.* [10] relying on HapCut’s tool extractHAIRS [2]. On a few real instances where SDhaP and H-PoP failed to run gracefully, we considered their assembled haplotypes to be fully unknown (all ·). We were unable to run HapTree [3] on any of our instances nor to communicate with its corresponding authors. To insure reproducibility of the results, we wrote a Snakemake pipeline [7] and deployed it on IBM Cloud™ using private single-core instances.

### Recall and precision

We measured precision and recall accounting for uncalled alleles in assembled haplotypes. That has been necessary because haplotype assembly tools omit to call some or all alleles at certain loci presumably due to insufficient sequencing data.

Given simulated haplotypes *X* of ploidy *p* over *n* alleles and assembled haplotypes *Y* with *µ* (*Y*) uncalled alleles, we computed the number of incorrectly called alleles as *δ* (*X, Y*) − *µ* (*Y*), where *δ* is phasing or haplotyping distance. Analogously, we computed the number of correctly called alleles as *np* − *δ* (*X, Y*) − *µ* (*Y*). We defined recall as the fraction of correctly called alleles over all alleles, which is equivalent to:

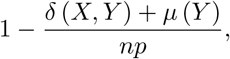

and precision as the fraction of correctly called alleles over all called alleles, which is equivalent to:

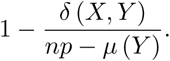

We defined genotyping recall and precision analogously. Both known genotypes *G* and induced genotypes *d*(*Y*) may contain unknown alleles. The number of called alleles is *np* − *µ*(*Y*). We computed the number of incorrectly called alleles as:

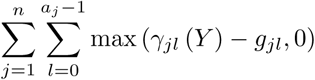

and the number of correctly called alleles as called minus incorrectly called alleles.

We defined recall and precision also in terms of MEC. Given the assembled haplotypes *Y*, we computed the number of fragments implied to be incorrect as EC (*Y, F*;*W*), those implied to be correct as:

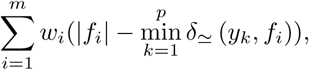

and the total number of fragments as 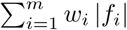.

### Results on simulated data

Table 1 shows mean precision and recall for each tool on the Illumina-like paired-end dataset with respect to haplotyping, phasing, genotyping and MEC, as well as mean runtime and memory footprint. Model *c*MFR obtained consistently the highest values in all precision and recall categories. In particular, *c*MFR obtained 8.4 % higher phasing recall than SDhaP. Model *a*MFR lost 1.1 to 1.4 % recall and precision in haplotyping and phasing. Note that HapCompass achieved perfect genotyping but it obtained the lowest recall and precision in haplotyping, phasing and MEC. We remark how MEC is in agreement with haplotyping and phasing when ranking the tools by recall and precision. Mean runtime for *c*MFR is 17.6 seconds while for *a*MFR it is 45.0 seconds. H-PoP and HapCompass achieved significantly lower runtimes compared to our MFR-based ILPs but their recall in all categories is equally lower. Memory footprint stayed within 120 MB for all tools except SDhaP that required 6 GB of main memory.

**Table 1:** Haplotype assembly results on simulated data. Mean values across all tested instances.

| Tool | Haplotyping |  | Phasing |  | Genotyping |  | MEC |  | Resources |  |
| --- | --- | --- | --- | --- | --- | --- | --- | --- | --- | --- |
|  | RecL.<br>[%] | Prec.<br>[%] | RecL.<br>[%] | Prec.<br>[%] | RecL.<br>[%] | Prec.<br>[%] | RecL.<br>[%] | Prec.<br>[%] | Time<br>[s] | Mem.<br>[MB] |
| cMFR | **98.0** | **98.1** | **95.8** | **95.8** | **100.0** | **100.0** | **99.9** | **99.9** | 17.6 | **48** |
| aMFR | 96.8 | 97.0 | 94.5 | 94.7 | 99.7 | 99.7 | 99.8 | **99.9** | 45.0 | **48** |
| H-PoP | 87.2 | 92.4 | 85.1 | 90.3 | 97.2 | 98.7 | 94.3 | 98.9 | **1.1** | 63 |
| SDhaP | 91.1 | 94.1 | 87.4 | 90.5 | 98.5 | 99.4 | 96.6 | 99.2 | 34.6 | 6309 |
| HapCompass | 86.8 | 86.9 | 81.8 | 82.1 | **100.0** | **100.0** | 92.6 | 92.8 | 4.7 | 112 |

Figure 3 (left) shows the distribution of haplotyping recall and precision values on the Illumina-like paired-end dataset. It is remarkable how model *c*MFR achieved 100.0 % median recall and precision. Figure S1 (left) shows recall and precision under phasing distance. Recall and precision values under phasing distance are lower with respect to haplotyping distance because phasing distance does not employ switch operations. Nonetheless, there is no significant change in the relative performances of haplotyping tools.

**Fig. 3:**
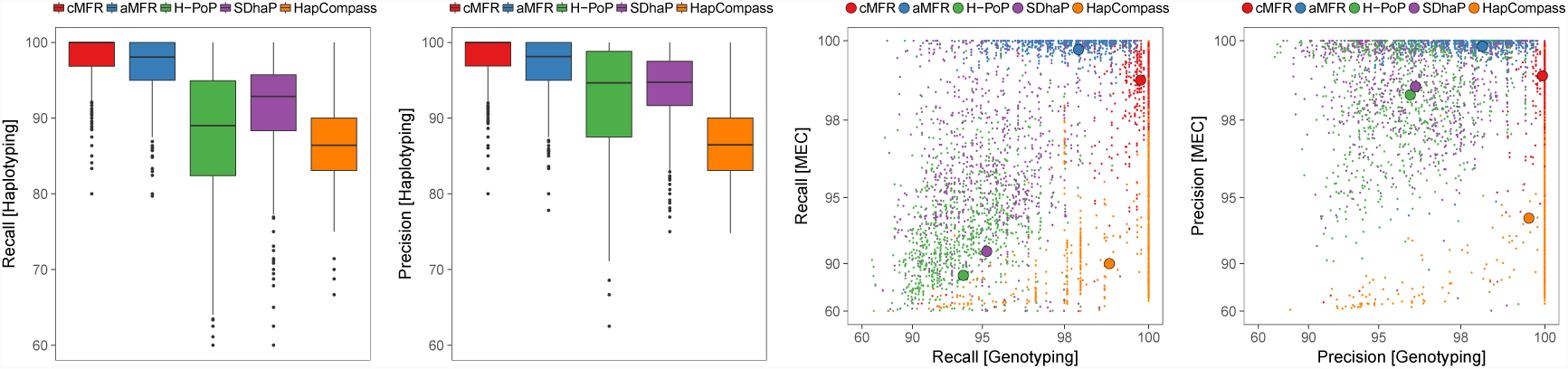
Haplotype assembly results. The two leftmost plots show haplotyping recall and precision on simulated data. The two rightmost plots show MEC versus genotyping on real data. Dots in the scatter plot denote results for individual instances, while circles denote mean values per tool.

Table S2 and Figure S2 (left) show results on the error-free paired-end dataset. Recall and precision values are in line with those on the Illumina-like paired-end dataset shown in Table 1. All tools show between 0.1 and 0.3 % improvement on the error-free dataset. That is in agreement with the 0.3 % sequencing error-rate used for Illumina-like reads simulation. None of the tools achieved perfect haplotyping recall or precision.

Table S3 and Figure S2 (right) show results on the mate-pair dataset. Model *c*MFR obtains 99.5 % mean recall and 99.7 % mean precision under haplotyping distance. That is an almost perfect assembly, 0.3 %. Conversely, none of the other assemblers shows a significant improvement on the mate-pair dataset with respect to the paired-end datasets.

### Results on real data

We computed genotyping recall and precision by comparing the genotypes induced by the assembled haplotypes to the genotypes previously computed by the variant caller. In addition, we computed precision and recall values for MEC as this can be done without knowing the true haplotypes. Recall and precision under MEC correlate well with haplotyping and phasing, as seen in Table 1.

Table 2 shows mean precision and recall of each tool across all real instances with respect to genotyping and MEC, as well as mean runtime and memory footprint. MEC precision for all tools is comparable to what observed on simulated data, while MEC recall on real data drops significantly for H-PoP and SDhap. Figure 3 (right) shows the distribution of MEC versus genotyping recall and precision on all real instances. As expected, *c*MFR and *a*MFR values are concentrated on the top right corner, with *c*MFR lying on (or very close to) the 100 % MEC line and *a*MFR lying on the 100 % genotyping line.

**Table 2:**
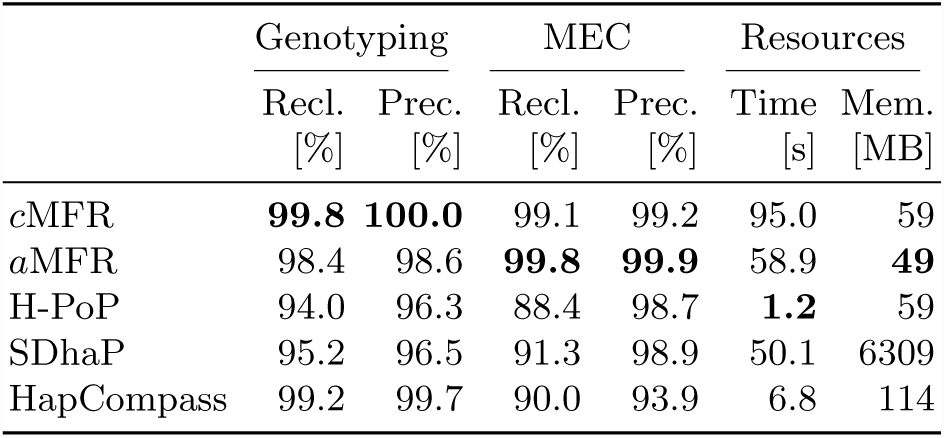
Haplotype assembly results on real data. Mean values across all tested instances.

## 4 Discussion

Motivated by practical limitations in breeding autotetraploid potato cultivars, we investigated methods for haplotype assembly of polyploids. We designed haplotype assembly models that are based on MFR, incorporate sequencing coverage using genotypes and are solved to optimality using integer linear programming. In addition, we proposed haplotyping distance to overcome the limitations of existing methods in comparing polyploid genomes. We applied haplotyping distance to evaluate the accuracy of haplotype assemblies.

Results on simulated data showed that our methods achieve a sensible improvement over the state of the art. Results on real data confirmed the relative improvements of our methods. Existing methods do not attain our performances either because of inadequate modeling, or because of heuristics failing to find optimal solutions, or because of implementation issues.

On similar Illumina short sequencing reads, we do not expect a MEC-based model to provide significant improvements over MFR. In fact, precision and recall results on error-free sequencing data indicate that residual errors in the assemblies are due to insufficient data (i.e., too short fragments) rather than erroneous sequencing data. Results on mate-pair data show that longer fragments boost assembly accuracy and thus support our hypothesis. Conversely, we expect long noisy reads to be challenging for MFR-based models. We did not investigate this latter hypothesis because we have no access to such data.

Our simulation does not faithfully reproduce all possible artifacts that may arise along high-throughput sequencing pipelines and cumulate along sample preparation, base calling, read mapping, variant calling and genotyping steps. In addition, our simulation does not account for the presence of copy number variations (CNVs) or complex structural variations in the genomic regions of interest. While none of the existing methods model CNVs, our ILPs are flexible enough to take in consideration known CNVs.

## A Supplementary results

**Table S1:**
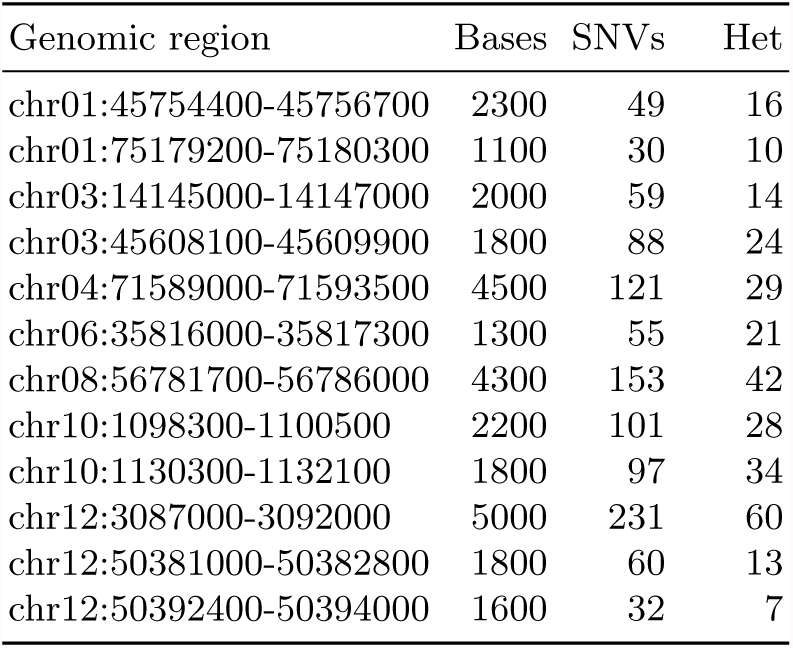
Genomic regions selected from the sequencing panel of Uitdewilligen *et al.* [15]. SNVs indicates the total number of SNP or Indel loci across the population, while Het indicates the median number of heterozygous loci per sample.

| Genomic region | Bases | SNVs | Het |
| --- | --- | --- | --- |
| chr01:45754400-45756700 | 2300 | 49 | 16 |
| chr01:75179200-75180300 | 1100 | 30 | 10 |
| chr03:14145000-14147000 | 2000 | 59 | 14 |
| chr03:45608100-45609900 | 1800 | 88 | 24 |
| chr04:71589000-71593500 | 4500 | 121 | 29 |
| chr06:35816000-35817300 | 1300 | 55 | 21 |
| chr08:56781700-56786000 | 4300 | 153 | 42 |
| chr10:1098300-1100500 | 2200 | 101 | 28 |
| chr10:1130300-1132100 | 1800 | 97 | 34 |
| chr12:3087000-3092000 | 5000 | 231 | 60 |
| chr12:50381000-50382800 | 1800 | 60 | 13 |
| chr12:50392400-50394000 | 1600 | 32 | 7 |

**Table S2:** Results on simulated error-free paired-end data. Mean values across all tested instances.

| Tool | Haplotyping |  | Phasing |  | Genotyping |  | MEC |  | Resources |  |
| --- | --- | --- | --- | --- | --- | --- | --- | --- | --- | --- |
|  | Recl.<br>[%] | Prec.<br>[%] | Recl.<br>[%] | Prec.<br>[%] | Recl.<br>[%] | Prec.<br>[%] | Recl.<br>[%] | Prec.<br>[%] | Time<br>[s] | Mem.<br>[MB] |
| cMFR | **98.2** | **98.2** | **96.0** | **96.0** | **100.0** | **100.0** | **100.0** | **100.0** | 8.8 | **45** |
| aMFR | 97.8 | 97.9 | 95.6 | 95.7 | 99.9 | 99.9 | **100.0** | **100.0** | 13.2 | **45** |
| H-PoP | 87.1 | 92.3 | 85.0 | 90.3 | 97.2 | 98.7 | 94.3 | 99.1 | **1.1** | 63 |
| SDhaP | 91.5 | 94.4 | 87.6 | 90.7 | 98.6 | 99.4 | 96.7 | 99.4 | 34.9 | 6309 |
| HapCompass | 86.8 | 86.9 | 81.9 | 82.2 | **100.0** | **100.0** | 92.6 | 92.8 | 4.7 | 112 |

**Table S3:** Results on simulated mate-pair data. Mean values across all tested instances.

| Tool | Haplotyping |  | Phasing |  | Genotyping |  | MEC |  | Resources |  |
| --- | --- | --- | --- | --- | --- | --- | --- | --- | --- | --- |
|  | Recl.<br>[%] | Prec.<br>[%] | Recl.<br>[%] | Prec.<br>[%] | Recl.<br>[%] | Prec.<br>[%] | Recl.<br>[%] | Prec.<br>[%] | Time<br>[s] | Mem.<br>[MB] |
| cMFR | **99.5** | **99.7** | **99.4** | **99.6** | 99.9 | **100.0** | **99.7** | **99.8** | 35.8 | 53 |
| aMFR | 98.1 | 98.5 | 98.1 | 98.4 | 99.6 | 99.7 | 99.6 | **99.8** | 57.4 | **51** |
| H-PoP | 88.1 | 91.8 | 86.9 | 90.6 | 97.4 | 98.5 | 93.4 | 97.6 | **1.2** | 62 |
| SDhaP | 90.2 | 91.6 | 87.4 | 88.9 | 98.5 | 98.9 | 95.9 | 97.8 | 35.0 | 6309 |
| HapCompass | 86.5 | 86.3 | 81.9 | 81.9 | **100.0** | **100.0** | 92.9 | 92.9 | 10.2 | 119 |

**Fig. S1:**
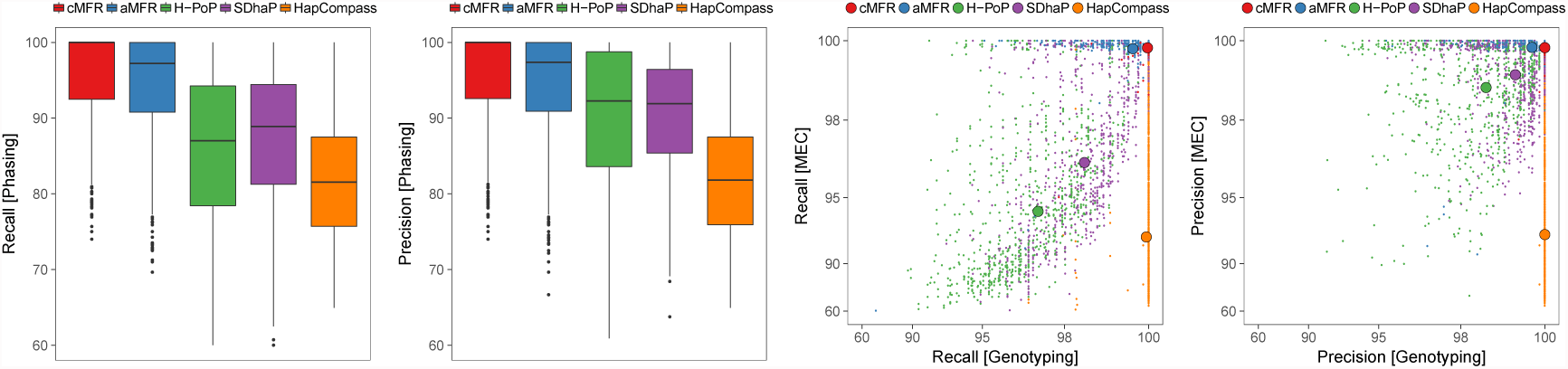
Results on simulated Illumina-like paired-end data. The two leftmost plots show phasing recall and precision. The two rightmost plots show MEC versus genotyping recall and precision. Dots in the scatter plot denote results for individual instances, while circles denote mean values per tool.

**Fig. S2:**
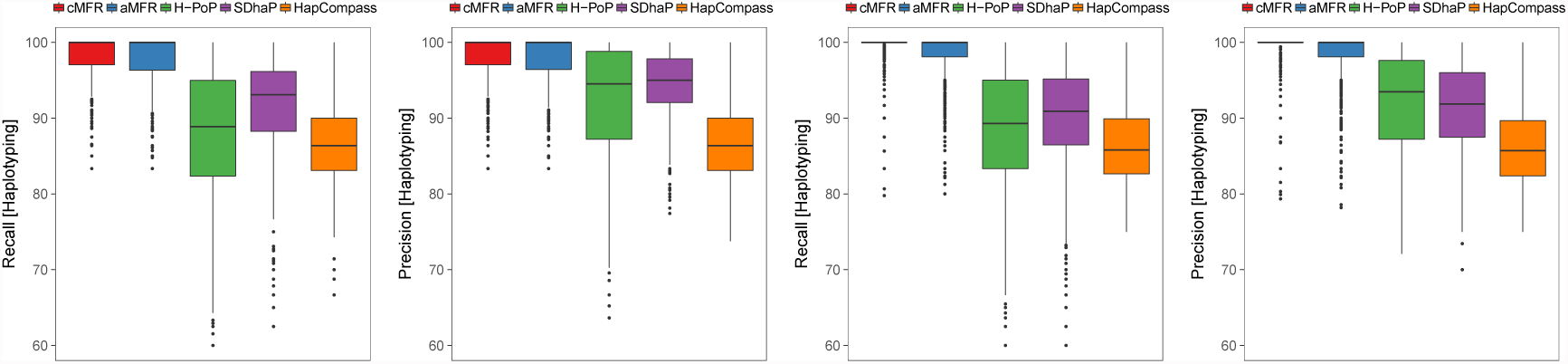
Haplotyping recall and precision on simulated data. The two leftmost plots show results on error-free paired-end data, while the two rightmost plots show results on mate-pair data.

